# Stability of excitatory structural connectivity predicts the probability of CA1 pyramidal neurons to become engram neurons

**DOI:** 10.1101/759225

**Authors:** Tim P. Castello-Waldow, Ghabiba Weston, Alireza Chenani, Yonatan Loewenstein, Alon Chen, Alessio Attardo

## Abstract

Neurons undergoing activity-dependent plasticity represent experience and are functional for learning and recall thus they are considered cellular engrams of memory. Although increase in excitability and stability of structural synaptic connectivity have been implicated in the formation and persistance of engrams, the mechanisms bringing engrams into existence are still largely unknown. To investigate this issue, we tracked the dynamics of structural excitatory synaptic connectivity of hippocampal CA1 pyramidal neurons over two weeks using deep-brain two-photon imaging in live mice. We found that neurons that will prospectively become part of an engram display higher stability of connectivity than neurons that will not. A novel experience significantly stabilizes the connectivity of non-engram neurons. Finally, the density and survival of dendritic spines negatively correlates to freezing to the context but not to the tone in a trace fear conditioning learning paradigm.

## INTRODUCTION

Ensembles of neurons undergoing coordinated activity-dependent plasticity (ADP) not only represent experience but are also functional for learning and memory recall (Barth, 2007; Cowansage et al., 2014; de Sousa et al., 2019; Denny et al., 2014; Guzowski et al., 2005; Kawashima et al., 2013; Liu et al., 2012; Marrone et al., 2008; Ramirez et al., 2013; Reijmers et al., 2007), thus they are largely believed to be cellular memory engrams (Josselyn et al., 2015; Mayford and Reijmers, 2015; Poo et al., 2016; Tonegawa et al., 2015; 2018). However, the cellular mechanisms that bring cellular engrams into existence and the cellular changes that engram cells undergo upon experience are largely unknown.

The transcription factor CREB has been implicated in memory formation (Kida et al., 2002; Restivo et al., 2009; Sekeres et al., 2010) and can bias neurons to become part of engrams by increasing their excitability (Dong et al., 2006; Han et al., 2007; 2009; Lopez de Armentia et al., 2007; Rogerson et al., 2016; Zhou et al., 2009). Increased excitability might be important for memory encoding and recall (Ballarini et al., 2009; Moncada et al., 2015; Pignatelli et al., 2019; Takeuchi et al., 2016). Moreover, neurons that have undergone ADP and engram neurons are more excitable and show higher activity levels (Barth et al., 2004; Ghandour et al., 2019; Kawashima et al., 2013; Yassin et al., 2010). While the level of excitability plays an important role in the selection and function of engram neurons, it is still unclear when and how prospective engram neurons become more excitable. To solve this issue it is necessary to study prospective engram neurons before they become part of an engram.

In the hippocampus, ensembles of neurons representing experience and undergoing ADP turn over with time (Attardo et al., 2018; Clopath et al., 2017; Hainmueller and Bartos, 2018; Mankin et al., 2012; Rubin et al., 2015; Ziv et al., 2013). Structural synaptic connectivity might underlie such turnover limiting the temporal window in which memories could be stored in the hippocampus (Attardo et al., 2018; 2015). Thus dynamics of structural synaptic connectivity in the hippocampus could have a direct relationship with hippocampal learning and recall - comparable to the neocortex (Hayashi-Takagi et al., 2015; Xu et al., 2009; G. Yang et al., 2009; Y. Yang et al., 2016). However, there is no clear correlation between hippocampal structural synaptic dynamics and hippocampal-dependent learning and recall.

To tackle these questions we used deep-brain two-photon (2P) time-lapse imaging (Ulivi et al., 2019) in transgenic mice in which neurons expressing the Immediate-Early Gene (IEG) Arc could be permanently marked during a specific time window (Guenthner et al., 2013). This enabled us to track the dynamics of structural excitatory synaptic connectivity of hippocampal CA1 engram and non-engram neurons both prospectively and retrospectively over a period of two weeks and to correlate such dynamics to hippocampal-dependent memory recall in a trace fear conditioning learning task, all in the same subjects.

## RESULTS

### Using the promoter of the Immediate Early Gene Arc to label CA1 engram neurons

To label neurons that became part of an engram during a defined time window, we used the Arc-Cre^ERT2^; Ai9 double knock-in mouse line (Guenthner et al., 2013), in which the IEG Arc drives expression of a Cre^ERT2^ recombinase, leading to tamoxifen (TAM)-dependent expression of tdTomato.

To test whether we could detect neurons that became part of an engram during exploration of an enriched environment (EE) in the dorsal CA1, we injected different groups of Arc-Cre^ERT2^; Ai9 mice with increasing doses of TAM (75 mg/kg, 150 mg/kg and 300 mg/kg). For each concentration of TAM one group explored an EE (**Fig. S1A**) for 2h, while the other group remained in its home cage (HC). All doses of TAM led to tdTomato expression (**Fig. S1B**), but a single 75 mg/kg TAM injection yielded a significant two-fold increase in the number of tdTomato + neurons after EE (**Fig. S1C, D**; * 2-way ANOVA corrected for multiple comparisons, **Table S1** for p values).

### Prospective and retrospective tracking of dendritic spines in CA1 engram neurons

To visualize dendritic spines in CA1 engram neurons, we obtained triple transgenic mice (Arc-Cre^ERT2^; Ai9; Thy1-eGFP) in which a sparse, random, subpopulation of CA1 pyramidal neurons (PNs) expressed cytoplasmic eGFP (Feng et al., 2000) in addition to activity-dependent tdTomato expression. In these mice, we tracked dendritic spines in the dorsal CA1 over two weeks using deep-brain 2P optical imaging (Ulivi et al., 2019) (Fig. 1A). This allowed us to longitudinally investigate the synaptic dynamics of CA1 engram and non-engram neurons within the same subjects, prior to and after EE.

**Figure 1.**
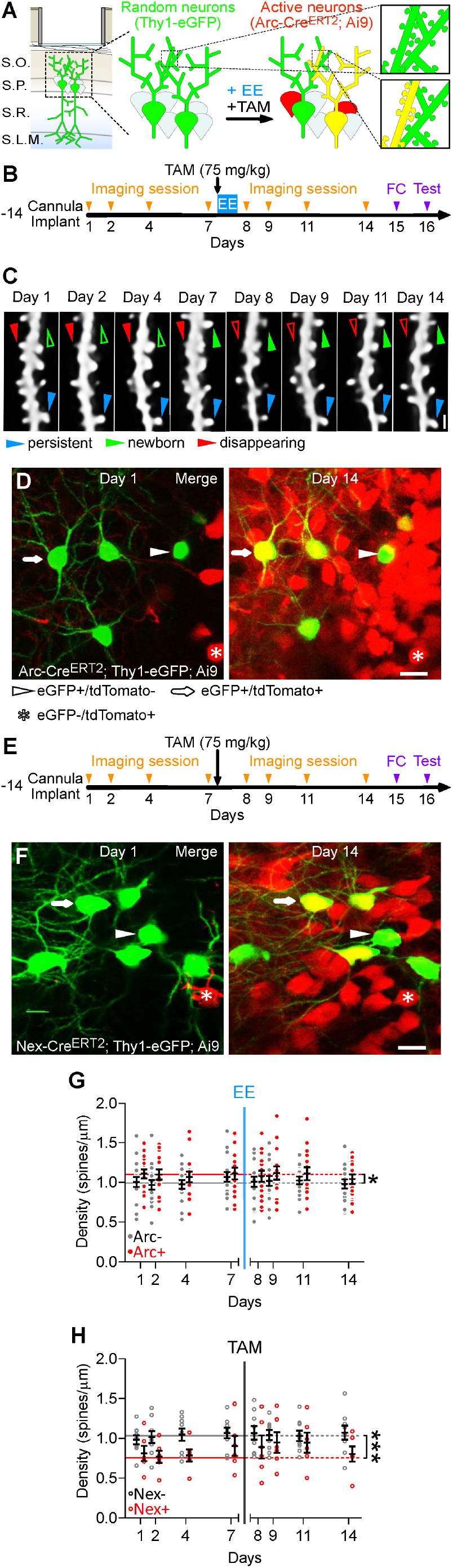
Retrospective and prospective tracking of dendritic spine dynamics of CA1 engram neurons in live mice. **(A)** Schematic description of the preparation and of the experiment‘s rationale. S.O., Stratum Oriens, S.P., Stratum Pyramidale; S.R., Stratum Radiatum; S.L.M., Stratum Lacunosum-Molecolare. EE, exposure to an Enriched Environment; TAM, Tamoxifen injection. Green, eGFP + neurons; red, tdTomato + neurons; yellow, eGFP / tdTomato double + neurons. **(B)** Experimental timeline. FC, trace fear conditioning memory task. **(C)** *In vivo* two-photon (2P) time-lapse of the same dendritic branch and spines over 14 days. Maximum intensity projections (MIPs) of up to 15 z-planes (Z step, 1 µm). Scale bar, 1 µm. **(D)** 2P images (MIPs of up to 5 z-planes, 2 µm z-step) of the dorsal CA1 of Arc-Cre^ERT2^; Ai9; Thy1-eGFP live mice. Scale bar, 20 µm. **(E)** Nex-Cre^ERT2^; Ai9; Thy1-eGFP; mice underwent the same experimental procedure as in A and B but were never exposed to EE. **(F)** 2P images (MIPs of up to 5 z-planes, 2 µm z-step) of the dorsal CA1 of Nex-Cre^ERT2^; Ai9; Thy1-eGFP live mice. Scale bar, 20 µm. (**G, H**) Spine densities of (G) ArcTom +, ArcTom -and (H) NexTom + and NexTom - neurons. Data points are mean density per neuron (**Table S2** for number of dendrites per neuron). Red: prospective or actual td-Tomato + neurons; grey: prospective or actual td-Tomato - neurons; solid circles: Arc-Cre^ERT2^; Ai9; Thy1-eGFP mice; empty circles: Nex-Cre^ERT2^; Ai9 Thy1-eGFP mice; black bars: means ± S.E.M; horizontal lines: median baseline densities per each group. **Table S1** for p values. Blue vertical line, time point of EE+TAM injection; grey vertical line, time point of TAM injection.

We imaged the dendritic spines of basal dendrites of CA1 PNs in seven Arc-Cre^ERT2^; Ai9; Thy1-eGFP mice for one week (**Table S2** for number of neurons and dendrites imaged). On the evening of day 7, we injected TAM and moved the mice to an EE. We returned the mice to their HC in the morning of day 8 and imaged them for one more week (Fig. 1B, C). 56.8% of the eGFP + neurons we tracked showed Arc-driven tdTomato expression (Fig. 1D, arrows), which we will refer to as Arc-tdTomato (ArcTom) + neurons. We excluded any neurons expressing tdTomato since the beginning of imaging from our analysis (Fig. 1D, asterisks).

To control for potential effects of repeated imaging, TAM injection, tdTomato expression and scoring variability we performed the same experiment -without exposure to EE (Fig. 1E) - on Nex-Cre^ERT2^; Ai9; Thy1-eGFP transgenic mice in which the Cre^ERT2^ recombinase is expressed in a subset of CA1 PNs independently from neuronal activity (Agarwal et al., 2012) (**Fig. S1E, F**). Approximately, 52% of the eGFP + neurons we tracked became tdTomato + (Fig. 1F, arrows) and we will refer to these as Nex-tdTomato (NexTom) + neurons. We excluded any neurons expressing tdTomato since the beginning of imaging from our analysis (Fig. 1F, asterisks)

The orders of the dendrites of ArcTom + and ArcTom - neurons we analyzed were similar to each other, likewise for NexTom + and NexTom - neurons (**Fig. S1G**). Spine densities of prospective ArcTom + and ArcTom - neurons and of NexTom + and NexTom - neurons were different during baseline [Mean densities (spines / μm): ArcTom +: 1.09, ArcTom -: 1.00, NexTom +: 0.86, NexTom -: 1.04; * and *** pairwise Mann-Whitney test, **Table S1** for p values), possibly reflecting stochastic differences between subpopulations of neurons. However, spine densities were stable after TAM injection with or without exposure to EE (Fig. 1G, H; one-sample Wilcoxon test each day’s data points against median baseline densities, **Table S1** for pvalues), thus demonstrating that repeated imaging, TAM injection and exposure to EE do not grossly affect the number of dendritic spines.

### Stability of excitatory synaptic connectivity predicts the probability of CA1 PNs to become engram neurons

Average stability of structural connectivity (Fig. 1G, H) can arise from slower or faster dendritic spines’ temporal dynamics. Neurons with slower dynamics of spines (i.e., small gains and small losses) are more stably connected with the same presynaptic partners, while neurons with faster dynamics (i.e., high gains and high losses) are less stably connected with their pre-synaptic partners. To distinguish between these two possibilities, we analyzed the temporal dynamics of gain and loss of dendritic spines.

Turnover rate was lower in prospective ArcTom + neurons than in ArcTom - neurons already before EE (Fig. 2A; * and ** Shuffle test, **S.T.A.R. methods** for details, **Table S1** for p values). Low turnover derived from the combination of lower gain and higher survival rates (Fig. 2B, C; * and ** Shuffle test, **S.T.A.R. methods** for details, **Table S1** for p values). After EE, gain rate decreased while survival rate increased in ArcTom - neurons thus leading to a decrease in the turnover rate (Fig. 2A, B, C; **\*, \*\*** and **\*\*\*** Wilcoxon test, **S.T.A.R. methods** for details, **Table S1** for p values). Thus ArcTom + and ArcTom - neurons became statistically indistinguishable (Fig. 2A, B, C; Shuffle test, **S.T.A.R. methods** for details, **Table S1** for p values).

**Figure 2.**
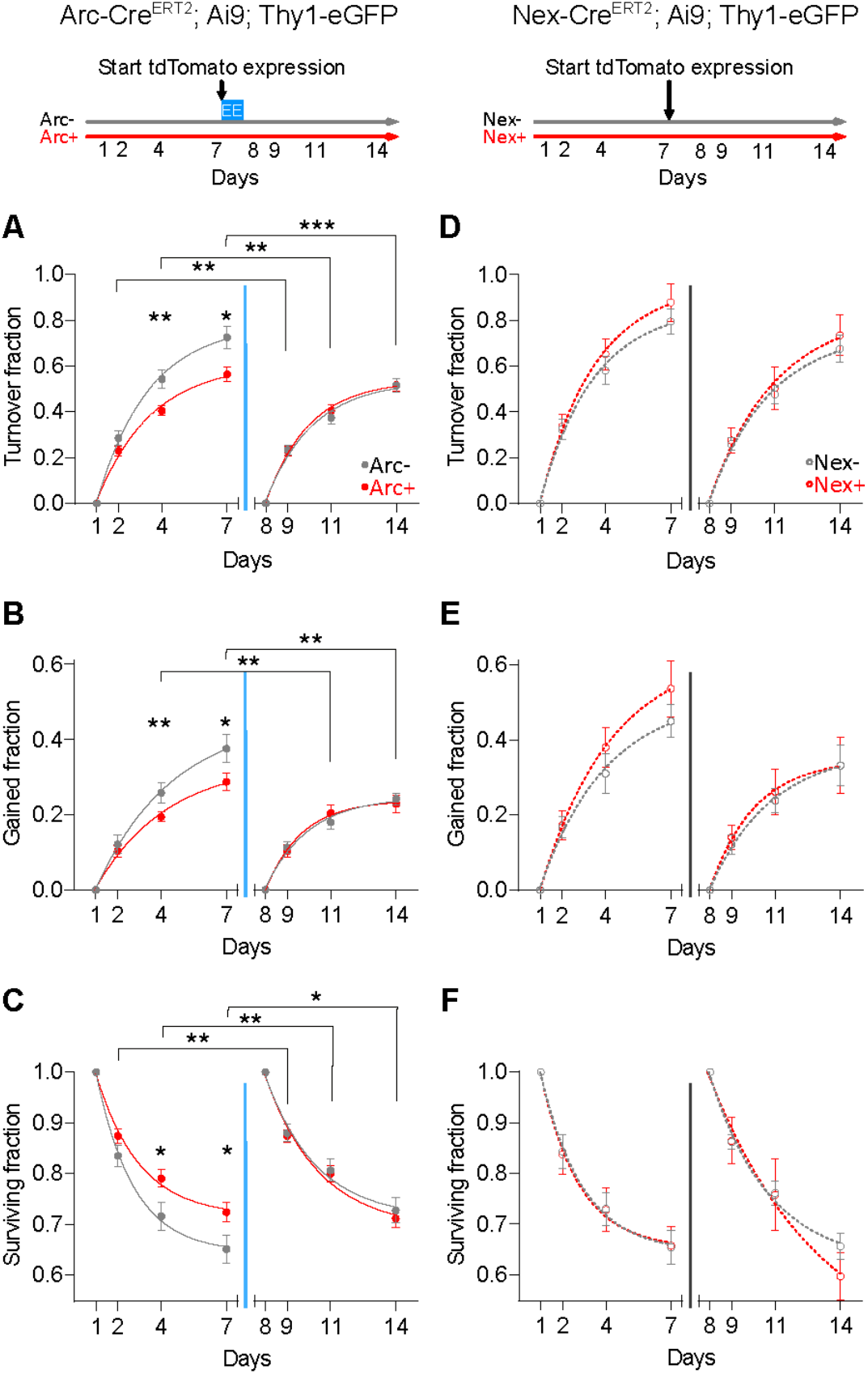
Stability of dendritic spines predicts the probability of CA1 PNs to become engram neurons. (**A, B**) The turnover fraction (A) and the fraction of dendritic spines gained (B) of prospective ArcTom + neurons was lower than prospective ArcTom - neurons prior to EE. The turnover and gain of ArcTom - but not ArcTom + neurons decreased after EE. (**C**) The survival fraction of prospective ArcTom + neurons was higher than prospective ArcTom - neurons prior to EE. Survival of dendritic spines of ArcTom - but not ArcTom + neurons increased after EE. (**D - F**) The turnover fraction (D), fraction of dendritic spines gained (E) and the survival fraction (F) of NexTom - and NexTom + neurons were not different during baseline and were not affected by TAM injection. (**A** - **F**) Data points are number of spines turned over (A, D), gained (B, E) or survived (C, F) normalized to day 1 or 8. Red: prospective or actual td-Tomato + neurons; grey: prospective or actual td-Tomato – neurons. Solid circles: Arc-Cre^ERT2^; Ai9; Thy1-eGFP mice; empty circles: Nex-Cre^ERT2^; Ai9; Thy1-eGFP mice. Curves: single exponential fits to the data: blue vertical line: time point of EE exposure +TAM injection; grey vertical line, time point of TAM injection. See **Table S1** for p values and **S.T.A.R. methods** for defintion of the error bars.

These results held true for the most part also when we compared turnover, gain and loss pooled over the entire baseline and after EE periods (**Fig. S2; \*** and **\*\*** Mann-Whitney test corrected for multiple comparisons, **Table S1** for p values).

As differences in spine sizes between ArcTom + and ArcTom - neurons or through time might bias the detectability of spines and lead to apparent differences in gain and survival, we quantified the variation in size of stable spines of ArcTom - and ArcTom + neurons through time. We found no difference between the sizes of ArcTom - and ArcTom + neurons and no variation through time (**Fig. S2G;** Mann-Whitney test; 2-way ANOVA respectively **Table S1** for p values), thus excluding the possibility that changes in spine gain and loss rates could be due to increase in spine size.

Analyses of turnover rate, spine gain and loss rates did not show any difference between NexTom + and NexTom - neurons (Fig. 2D-F; Shuffle test, **S.T.A.R. methods** for details, and **Fig. S2D** - **F**; Mann-Whitney tests corrected for multiple comparisons, **Table S1** for p values).

Altogether, prospective engram neurons displayed higher excitatory structural synaptic stability even before their activation, and exposure to EE increased the synaptic stability of non-engram neurons.

### Exposure to EE stabilizes dendritic spines on recurrent sites of CA1 non-engram neurons

Approximately 35% of dendritic spines disappeared and recurred again at the same site along the dendrite during the experiment (Fig. 3A, B), and recurrent sites were occupied by a spine more often than expected by chance (**Fig. S3A**, **B**; Probability mass test, **S.T.A.R. methods** for details, **Table S1** for p values). The positional jitter of recurrent spines was indistinguishable from the jitter of stable spines (Fig. 3C), with 90.1% of recurrent spines being located within 1 standard deviation of the distribution of the distances between the location on the dendrite at which they first appeared and the locations at which they recurred on following time points (2.7 µm, Fig. 3C, **S.T.A.R. methods** for details). As spines at recurrent sites are very likely to connect the same pre- and post-synaptic neurons (Stepanyants et al., 2002), we decided to investigate the stability of recurrent dendritic synaptic sites.

**Figure 3.**
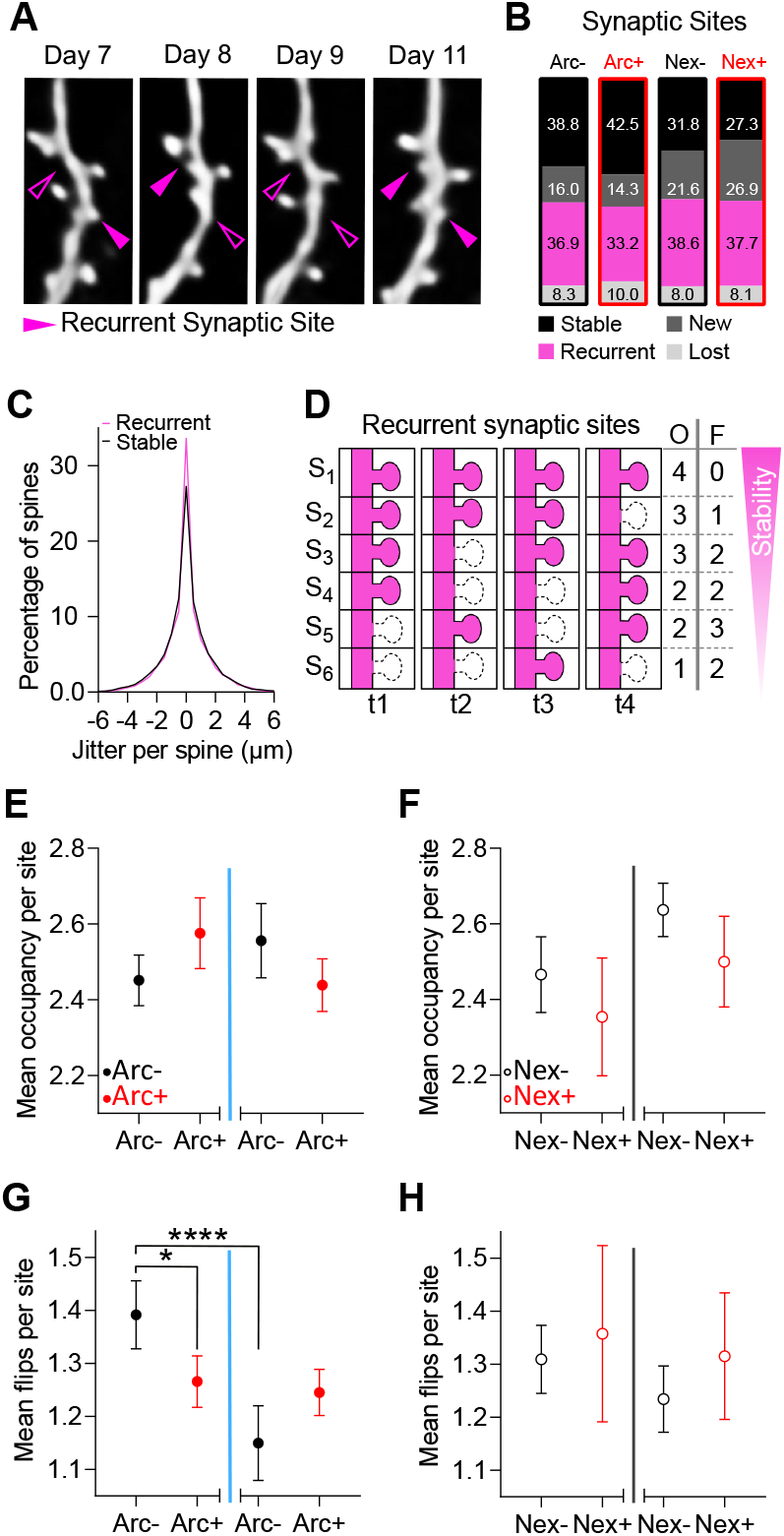
Exposure to EE stabilizes dendritic spines on recurrent sites of CA1 non-engram neurons. **(A)** *In vivo* 2-photon time-lapse of the same dendritic branch and spines over 5 days. Maximum intensity projections (MIPs) of up to 7 z-planes (Z step, 1 µm). Scale bar, 1 µm. **(B)** Percentages of stable, new, recurrent, and lost synaptic sites defined over the 14 imaging days. **(C)** The distributions of positional jitters for stable (black) and recurrent (magenta) dendritic spines were not different. **(D)** Schematic description of recurrent synaptic sites at consecutive time points displaying different numbers of time points (t) occupied (O) and flips (F). (**E, F**) Occupancy of recurrent sites of ArcTom -, ArcTom +, NexTom - and NexTom + neurons were not different during baseline and were not affected by exposure to EE or TAM injection. **(G)** The number of flips of recurrent sites was lower in ArcTom + than in ArcTom – neurons. The number of flips of recurrent sites of ArcTom - neurons decreased upon EE. **(H)** The number of flips of recurrent sites sites of NexTom - and NexTom + neurons were not different during baseline and were not affected by TAM injection. (**E - H**) Data points are mean number of time points occupied (E, F) or mean number of flips (G, H) during baseline, after exposure to EE or TAM injection of ArcTom - (solid circles, black), ArcTom + (solid circles, red), NexTom - (open circles, black) and NexTom + (open circles, red) neurons. **S.T.A.R. methods** for defintion of the error bars. (**C**, **E - H**) See **Table S1** for p values.

When we examined occupancy (Fig. 3D, O) of recurrent sites (Fig. 3, E, F and **Fig. S3C, F**) we found no signficant differences during baseline and after EE (Fig. 3E, F). However, when we quantified the number of transitions between occupied and not occupied states (or Flips, Fig. 3D, F; **Fig. S3G, J**), we found that recurrent sites of ArcTom + neurons showed fewer flips than recurrent sites of ArcTom - neurons before EE and that exposure to EE decreased the number of flips of ArcTom - neurons (Fig. 3G, * and **** Mann-Whitney tests corrected for multiple comparisons, **S.T.A.R. methods** for details, **Table S1** for p values). These results support both the increased synaptic stability of prospective engram neurons before EE and the increase in synaptic stability of non-engram neurons after EE.

### Density and survival of dendritic spines of CA1 PNs correlate with hippocampal-dependent memory

Next, we investigated the relationship between dendritic spine’s dynamics and memory recall in a hippocampal-dependent learning task. After *in vivo* imaging we trained the animals in a trace fear conditioning (FC) memory task and probed their freezing to the context and to the tone separately on the next day (Fig. 1C, E and Fig. 4A).

**Figure 4.**
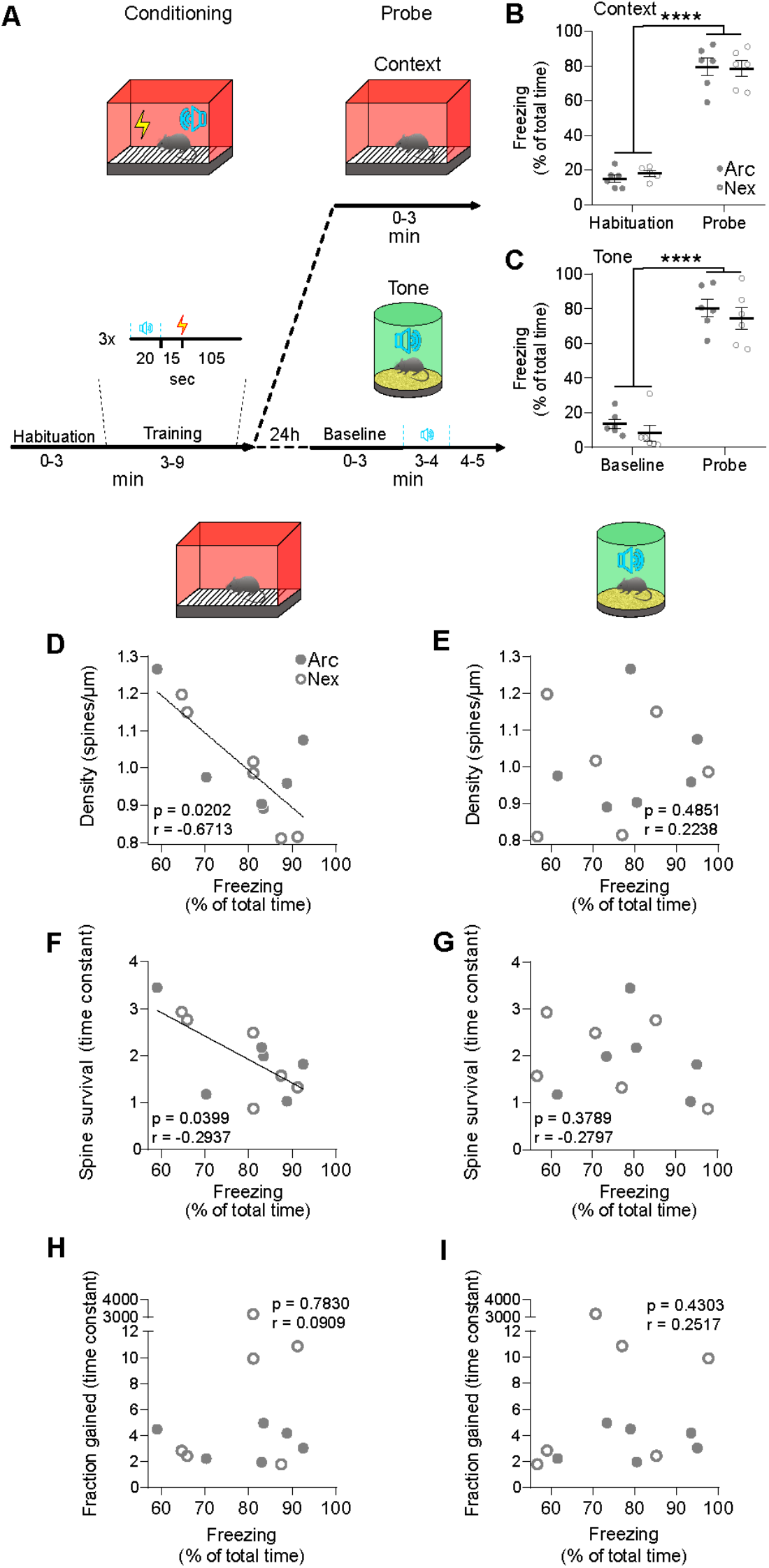
Density and survival of dendritic spines of CA1 PNs negatively correlate with freezing in a trace fear conditioning task. **(A)** Schematic description of the trace fear conditioning learning task. (**B, C**) Arc-Cre^ERT2^; Ai9; Thy1-eGFP (Arc, solid circles) and Nex-Cre^ERT2^; Ai9; Thy1-eGFP (Nex, open circles) mice displayed increased freezing during the context (B) and tone (C) probe trials. Circles are percentage freezing for each mouse during: 3 min habituation (day 15) and 3 min context probe (day 16)(B) or during 3 min baseline (day 16) and 4 min tone probe (day 16)(C); horizontal bars: means ± S.E.M. (**D** - **I**) Correlations between densities (D, E), time constants of the survival (F, G) and gain (H, I) rates of spines during baseline of Arc-Cre^ERT2^; Ai9; Thy1-eGFP (solid circles) and Nex-Cre^ERT2^; Ai9; Thy1-eGFP (open circles) and freezing to the context (D - F) and the tone (G - I). (**B** - **I**) See **Table S1** for p values.

Mice showed significant freezing to the context and to the tone (Fig. 4B, C; **** 2-way ANOVA corrected for multiple comparisons, **Table S1** for p values) demonstrating they had learned the association of the context and tone to the mild shock.

Average density and survival rates of spines of all neurons per mouse during baseline predicted the level of freezing to the context but failed to predict the level of freezing to the tone alone (Fig. 4D - G; * Spearman correlations, r_Density_ = −0.67, r_Survival_ = −0.29, r_Gain_ = −0.09, **Table S1** for p values). Average gain rates per mouse did not predict the level of freezing to the context or the tone (Fig. 4H - I; Spearman correlations, **Table S1** for p values). When we analyzed Arc-Cre^ERT2^; Ai9; Thy1-eGFP and Nex-Cre^ERT2^; Ai9; Thy1-eGFP mice separately after EE (or TAM alone), we found that the average density, the survival and the gain of spines did not predict freezing (**Fig. S4A - L**; Spearman correlations, **Table S1** for p values).

Altogether, these data suggest a relationship between baseline spine dynamics of CA1 PNs and hippocampal-dependent, but not hippocampal-independent learning and memory.

## DISCUSSION

### Prospective and retrospective tracking of dendritic spines in CA1 engram neurons

We investigated the connectivity of engram neurons not only prospectively but also retrospectively and compared the dynamics of structural connectivity of engram and non-engram neurons in the hippocampal CA1 of the same animals, thanks to the combination of 2P deep-brain imaging and genetic labelling of engram neurons (Guenthner et al., 2013).

While 2P imaging is widely used to track neuronal structure (Moyer and Zuo, 2018), it cannot resolve all spines in hippocampal CA1 (Attardo et al., 2015), leading to conflicting results (Attardo et al., 2015; Gu et al., 2014; Pfeiffer et al., 2018). To tackle this issue, we performed internally controlled experiments, by comparing spine dynamics of engram and non-engram neurons and spine dynamics of neurons before and after they became engram neurons, within the same subjects. In addition, we tracked synaptic dynamics of neurons that expressed the same fluorescent tag as engram neurons but in a non-activity-dependent fashion (Nex-Cre^ERT2^ mice (Agarwal et al., 2012)) to control for the noise generated by optical resolution, repeated imaging, manual scoring of spines, tdTomato expression and TAM injection. Thus, while we cannot provide an absolute measure of the synaptic stability of neurons, we could uncovered significant differences in synaptic stability.

We did not detect a significant change in spine density in the dorsal CA1 after 16 h of EE, in contrast to others who have reported density increases in the neocortex and hippocampus (Jung and Herms, 2012; Moser et al., 1994; Rampon et al., 2000) or density decreases in CA1 PNs expressing Arc (Kitanishi et al., 2009) upon EE. This discrepancy might be due to the different duration of the EE, to the time after EE at which spine density was quantified or to the statistical fluctuations between different groups of animals.

### Prospective CA1 engram neurons display higher structural synaptic connectivity

A key finding of our work is that, despite stationary average structural connectivity, CA1 engram neurons stay connected for longer durations of time with their pre-synaptic partners during the week prior to becoming active (Fig. 2A - C).

Especially in a brain region with high synaptic turnover such as hippocampal CA1 (Attardo et al., 2015; Pfeiffer et al., 2018), a bias towards structural synaptic stability might result in increased functional connectivity, which in turn would render a subset of neurons more receptive to memory allocation (Epsztein et al., 2011; Restivo et al., 2009; Sekeres et al., 2010). Moreover, increased synaptic transmission could lead to local plasticity (Losonczy and Magee, 2006; Losonczy et al., 2008) and prime CA1 PNs to become more active (Bittner et al., 2017; Sheffield et al., 2017). Further experiments should clarify the extent to which high structural stability leads to increased neuronal excitability.

As activity patterns of pyramidal neurons are more stable in CA3 than CA1 (Mankin et al., 2012), pyramidal CA1 PNs with higher connectivity to CA3 neurons could potentially be a subpopulation of active neurons (Ghandour et al., 2019) biased towards temporal stability (Karlsson and L. M. Frank, 2008). Still, another prominent projection to CA1 - area CA2 - displays unstable temporal activity patterns (Mankin et al., 2015) and further experiments are needed to clarify the differential influence of CA3 and CA2 regions on CA1 activity patterns.

### Synaptic stabilization of non-engram neurons

In our study we found stabilization of connectivity upon exploration of a novel environment, which is in line with previous studies described stabilization of connectivity of PNs upon learning in the neocortex (Xu et al., 2009; G. Yang et al., 2009; Y. Yang et al., 2016). However, while these studies could not distinguish between engram and non-engram neurons, we found that stabilization of connectivity occurred predominantly in CA1 non-engram neurons. One possible explanation for our finding is that engram neurons produce factors that stabilize connectivity but these factors affect mostly non-engram neurons, possibly due to a ceiling effect whereby engram neurons are already synaptically stable and cannot be stabilized any further. Another intriguing possibility is that such factors could specifically target neurons that do not express ADP-related genes. For instance, it has been shown that the protein Arc can mediate the transfer of Arc mRNA which in turn can be translated in neighboring neurons (Pastuzyn et al., 2018). This extracellular signaling might inform non-engram neurons of the status of their neighbors undergoing ADP and be functional for system consolidation (Pastuzyn et al., 2018).

### Higher density and survival of CA1 dendritic spines during baseline predict lower freezing to the context but not to the tone

We previously suggested that hippocampal dendritic spine dynamics reflect the function of the hippocampus in learning and memory (Attardo et al., 2015). Here we show that the density and the survival of hippocampal dendritic spines predict hippocampal-dependent recall (freezing to the context) but not hippocampal-independent recall (freezing to the tone). This finding suggests that also in the hippocampus - as in the neocortex (Hayashi-Takagi et al., 2015; Xu et al., 2009; G. Yang et al., 2009; Y. Yang et al., 2016) - structural synaptic dynamics can support brain area-specific cognitive functions.

Interestingly, animals with higher baseline density or stability of CA1 dendritic spines showed lower levels of freezing, similarly to what has been shown in retrosplenial cortex (A. C. Frank et al., 2018). This suggests that the ability of neurons to sample more pre-synaptic partners *per* unit time might support more effective learning. However, lower synaptic stability is also associated with cognitive impairments (Contractor et al., 2015; Cruz-Martín et al., 2010; Goel and Portera-Cailliau, 2019) and it will be important to elucidate the molecular and circuit mechanisms that mediate the effects of synaptic stability on cognition.

## ACKNOWLEDGEMENTS

We would like to thank Dr. Ju Lu for writing the first version of the Matlab GUI we use to track dendritic spines and prof. Luqun Luo for early access to the TRAP mice. We are also very grateful to Prof. Carsten Wotjak and Prof. Anton Sirota for their helpful comments. Y. L. is supported by the Israel Science Foundation (ISF), the Deutsche Forschungsgemeinschaft (DFG) and the Gatsby Charitable Foundation; A. C. is supported by an FP7 Grant from the ERC, the ERANET and I-CORE programs, the Israeli Ministry of Health, the BMBF, the Nella and Leon Benoziyo Center for Neurological Diseases, the Henry Chanoch Krenter Institute for Biomedical Imaging and Genomics, The ISF, the Perlman Family, the Adelis, Marc Besen, Pratt and Irving I. Moskowitz foundations and by Roberto and Renata Ruhman, Bruno and Simone Lich; A. A. is supported by the Max Planck Society, DFG and Schram Foundation.

## AUTHOR CONTRIBUTIONS

Conceptualization, Methodology and Writing – Original draft, T. C-W. and A. A.; Investigation, T. C-W.; Software, T. C-W. and G. W.; Formal analysis, T. C-W, A. Chenani. and Y. L.; Funding acquisition, A. Chen and A.A.; Supervision, Project Administration, A. A.

## S.T.A.R. METHODS

### Contact for Reagent and Resource Sharing

Further information and requests for reagents may be directed to and will be fulfilled by the Lead Contact, Dr. Alessio Attardo. All published reagents can be shared on an unrestricted basis.

### Experimental Model and Subject Details

Double heterozygous Arc-Cre^ERT2^ (+/−); Ai9 (+/−) mice were obtained by crossing Arc-Cre^ERT2^ (+/−) x Ai9 (+/+) mice. Crossing Thy1-eGFP (+/+) x Ai9 (+/+) mice yielded double homozygous Thy1-eGFP (+/+); Ai9 (+/+) mice which were further crossed with Arc-Cre^ERT2^ (+/−) or Nex-Cre^ERT2^ (+/+) mice to generate triple-transgenic Arc-Cre^ERT2^ (+/−); Ai9 (+/−); Thy1-eGFP (+/-) or Nex-Cre^ERT2^ (+/−); Ai9 (+/−); Thy1-eGFP (+/-) mice respectively. Mice were held on a 12-h light/dark cycle. Experiments were conducted during the 12-h light period. Littermates were group-housed, up to 5 mice per cage, with food and water *ad libitum*. All experimental procedures were conducted with 12 weeks old animals from either sex. All animal procedures conformed to the guidelines of the Max Planck Society and the Animal authority, Regierung Oberbayern (Regierung von Oberbayern – Veterinärwesen) and were in line with the Tierversuchslizenz - ROB-55.2Vet-2532.Vet_02-16-48.

## Method Details

### Labeling of neurons active during exploration of EE

Mice received an <u>intraperitoneal injection</u> of <u>tamoxifen</u> (75 mg/kg of body weight) right before being placed into the EE. Tamoxifen was dissolved in 5% of the final volume in 100% Ethanol and further diluted with corn oil to a final concentration of 10mg/ml. The solution was heated up to 37°C before injection. After exposure to the EE (2 h or 16 h) mice were transferred back into their HC. Enriched environments were created by connecting two rat-cages (37 cm x 60 cm) with an acrylic tunnel (20 cm x 15 cm) resulting in a total area 4440 cm^2^. Cages contained tunnels, wooden climbing sticks, wooden shelters, running wheels, seesaws, cotton pads, hair curlers, wooden blocks, swinging hammocks and toys which mice could open and which contained food pellets. Cages also contained a second level connected to the ground floor by a wooden ladder and consisting of a wooden board and climbing ropes allowing mice to reach the lid grit. Food was hidden in the bedding material and spread around the arena to encourage mice to explore the environment.

### Histology

We perfused mice intracardially with 1x phosphate-buffered saline (PBS) containing Heparin followed by 4 % paraformaldehyde (PFA) in PBS. We then dissected the brains and placed them in 4 % PFA in 1x PBS for 24 hours. Brains were then transferred to 30% sucrose in PBS for 48 hours. Brain slices (50 μm thick) were prepared with a vibratome (Thermo Scientific Microm HM 650V).

Slices were quenched with 150 mM Glycine in ddH20 for 15 minutes and permeabilized with 0.2 % Triton X-100 in PBS for 1 hour. Slices were incubated with DAPI (1:1000 diluted in PBS, Thermo Fisher) for 5 minutes washed with PBS and mounted onto slides with mounting medium (Vectashield).

### Cell Counting

To quantify the proportion of Arc- or Nex-tdTomato+ CA1 pyramidal neurons *ex vivo* we used a confocal microscope (Zeiss LSM 800) and acquired image stacks (319.28 µm^2^ single section area, 5 μm z-step, 8-10 focal planes) of 4 representative fields in the dorsal CA1 field per mouse using a 40x objective (Zeiss Plan-Apochromat 40x/1.4 Oil DIC (UV) VIS-IR). We acquired DAPI fluorescence (405 nm excitation and 465 nm emission wavelengths) to identify neuronal nuclei and tdTomato fluorescence (561 nm excitation and 618 nm emission wavelengths) to identify Arc- or Nex-tdTomato+ cells. We then manually counted DAPI+, tdTomato+ and double positive neurons using the plugin Cell Counter (Image J). When the same neuron was visible in more than one z-slice we only counted it once in the z-plane in which the diameter of the soma was the largest.

### Preparation of the imaging cannula

The imaging cannula was prepared as previously described (Ulivi A. et al., 2019). We cut a 1.6 mm-long and 3 mm diameter ring from stainless steel tubing and removed irregularities from the ring edge using a rotating cylindrical file and a dental drill. One edge of the metal ring was dipped into UV-curing optical adhesive (NOA81) and placed onto a 4 mm diameter glass coverslip (0.13 mm thick) before it was illuminated/cured with 365 nm light for 1 minute. 24 h later the excessive glass was filed away, using a rotating cylindrical file and a dental drill.

### Implantation of the imaging cannula

Surgeries were performed as previously described (Ulivi A. et al., 2019). To induce anesthesia mice were put into an anesthesia induction chamber which was flooded with 2.5% Isoflurane in pure O2. During the surgery, the anesthesia was held by 1.5% Isoflurane in pure O2 directly delivered to the mouse’ nostrils. The appropriate depth of anesthesia was confirmed by the absence of the toe pinch reflex. Meloxicam (1 mg/kg) and Vetalgin (200 mg/kg) were administered by subcutaneous injection and ophthalmic ointment (Bepanthen cream) was applied to protect the eyes. After hair removal, the skin was disinfected, and the skull - roughly from the frontal to the interparietal bone - was exposed. Non-rupture ear bars (David Kopf Instruments) were positioned to stabilize the skull, and a drop of Lidocaine (≈10 mg) was applied to the skull. A small craniotomy of the left frontal bone was performed using a microdrill with 0.5-mm burr. A 0.86 diameter stainless-steel screw was inserted and fixed by the application of Metabond. A 3 mm diameter craniotomy was performed using a trephine drill, the dura was removed, and the cortical matter was slowly ablated using a blunt needle connected to a vacuum pump until reaching the fibers of the corpus callosum. The imaging cannula was inserted into the craniotomy, applying slight pressure onto the tissue to stabilize the preparation, and the cannula was fixed and sealed to the skull using Metabond (Parkell). A custom head plate was positioned and fixed with dental cement. The following two days, mice received Meloxicam (1 mg/kg) once per day for postoperative analgesia.

### *In vivo* 2-photon imaging

Two to three weeks after implantation of the imaging cannula, mice were anesthetized with 2.5% Isoflurane and placed under the microscope (Bruker Ultima IV) onto a 37°C heating pad (CMA 450) while the head was fixed via a head plate holder. During imaging mice were kept under constant anesthesia (1.5% Isoflurane). To manually align the imaging cannula to the light path we used a 4x objective (Olympus Plan N 4x/0.10) and fluorescent light (X-Cite 120Q) to visualize the top and the bottom of the cannula. We adjusted the three separate rotational degrees of freedom of the animal’s head independently to align the axis of the cannula to the axis of imaging. We considered the cannula aligned when the top and bottom circular ends of the cannula were concentric. For 2P imaging the objective was changed to a 25x water immersion objective (Olympus XLPlan N 25x/1.00 SVMP). To excite eGFP and tdTomato we used a pulsed infrared laser tuned to 920 nm and 1040 nm respectively. On day 1 and 14, overview images of the eGFP and the tdTomato channel were acquired consecutively with 1x, 2x, and 5x digital zoom to relocate the dendrite of interest and to identify double positive neurons. To image dendritic spines we acquired z-stacks (48.18 µm^2^ single section area, 1 µm z-step, 5-60 z-steps, zoom 10x, 28.6-115.5 mW laser power at the sample) of the dendrite of interest using a resonant scanner, at each time point. We acquired each z-plane four times before moving to the next plane.

### Image post-processing

To compensate for motion artifacts, we split each of the stacks into four single stacks containing a single acquisition per each x plane. We then registered each stack in the x dimension by using the ImageJ plugins Turboreg and Stackreg (Philippe Thévenaz, GitHub) and deconvolved each stack separately using blind deconvolution (AutoquantX3). Finally, we registered the 4 corresponding z planes from each stack and averaged them using Image J. We aligned once more the resulting average stack through the z plane.

### Scoring of dendritic spines and identification of dendritic order

Dendritic spines were manually scored using a custom graphical-user-interface written in MATLAB (MathWorks) which concatenated all time points in a loop and hid the date in which each time point was acquired, thus the operator was blind to the experimental condition while counting. First, we traced dendrites in three dimensions with a node-connected line that permitted measuring the length of the dendritic segment. Then we marked a spine in one time point and moved to the next time point to identify the same spine. Initially, we marked 5 spines per dendrite present in all 8 timepoints which were used as stable fiduciary points for rigid-body registration across time. After registration, we counted all spines in each dendritic segment. Spines that disappeared were scored as lost spines on that day, while spines that were not present at the previous imaging timepoint were counted as new spines. New spines that appeared at a previously lost position were considered spines in a recurrent location.

Two-photon overview image stacks were used to identify if a dendrite belonged to a tdTomato+ neuron by manually tracing each dendritic segment back to its soma and to establish the dendritic order. Each basal dendrite originating from the soma was classified as a first-order dendrite. After the first branching point, both emerging dendrites were allocated as a second-order dendrite, independent of the size or the diameter of the two dendrites. Using this technique, the most distal identified dendritic segments belonged to the sixth order.

### Metrics used to quantify spine dynamics and size

In **Fig. S2**, fractional gain and fractional loss were defined as the number of spines gained or lost between each time point and the next one, normalized by the number of spines present in the first time point. Turnover rate was defined as the sum of the fractional gain and fractional loss. All values during baseline or aftger EE / TAM were pooled together and treated as a single distribution. To quantify the size of a spine we drew manually a region of interest (ROI) around a spine in a single z plane. We then calculated the sum of the fluorescence of all pixels included in the ROI of the spine - integrated density of fluorescence (IDF) - using ImageJ. The IDF thus accounted for the size of the ROI and the fluorescence of the ROI. The IDF of the spine was normalized to the IDF of a fixed-sized ROI of the dendritic shaft of the same z-plane below the spine.

In Fig. 2, fractional gain and fractional survival were defined as the number of spines gained or surviving between the first and each of the other time points, normalized by the number of spines present in the first time point (Baseline) or after EE/TAM. Fractional turnover rate was defined as a sum of the fractional gain and fractional loss (1 - surviving fraction).

In Fig. 3, The jitter of a spines’ location over time was defined as the distribution of the differences between the position of that spine on the linearized dendrite at the first time point it appeared and the positions of the same spine at all other time points in which it was detected. Recurrent sites were defined over the whole duration of the experiment (8 time points) and were characterized by a spine present at least at one time point, then lost and then reappearing at least once in a location that was visually indistinguishable from the first location it was detected in. The Occupancy (O) was defined as the number of time points the recurrent site contained a spine. A Flip (F) was defined as the appearance or the disappearance of a spine in a recurrent site.

In Fig. 4 and **Fig. S4**, the density of spines per mouse was calculated as the average of the densities of the dendrites of that mouse. To calculate the time constant tau of the turnover, gain and surviving rates per mouse we fitted exponential growth to plateau (for turnover and the growth) or single exponential decay (for survival) curves to the distributions of fractional turnover, gain or survival of the dendrites of that mouse.

### Trace Fear Conditioning

After two weeks of *in vivo* two-photon imaging, all mice were trained in a hippocampus-dependent trace fear conditioning task. On the training day (Day 15) mice were put into a square conditioning chamber (19 cm x 19 cm, black metal walls, stainless steel grid floor, white light illumination, and ethanol odor) (Panlab) which we defined as Context A. Following 3 minutes of habituation, mice received 3 pairings of a tone (80 dB, 9 kHz, 20 s duration, CS) and a mild electric footshock (0.75 mA, 1 s duration, US) with a trace of 15 s between the tone and the shock and an intra trial interval of 105 seconds. On day 16 mice were probed for their hippocampal-dependent and independent memory recall. To test hippocampal-dependent memory recall we placed mice into Context A for three minutes. Thirty minutes later, to test hippocampal-independent memory recall, mice were placed into a novel context (a round chamber of 15 cm diameter, transparent acrylic walls, bedding, white light illumination, and acetic acid odor) which we defined as Context B. Mice explored Context B for 4 minutes after which we administered the CS for one minute. During all exposures to Context A and B the position of the mouse was tracked automatically and the freezing response was recorded and quantified in real-time with ANY-maze (Stoelting). The amount of freezing was calculated as the percentage of total exploration time during which the mice were immobile. Immobility for more than 250 ms was scored as freezing.

### Quantification and Statistical Analysis

For statistics, we used the Mann-Whitney test (two-tailed), 1-sample Wilcoxon test, paired Wilcoxon, 2-way ANOVA, and Spearman correlation. For post hoc analysis for multiple comparisons, we used Šidák or Tukey’s test. Statistical analysis and plotting was done with Prism 8 (GraphPad) software.* p ≤ 0.05, ** p ≤ 0.01, *** p ≤ 0.001, **** p < 0.0001.

In **Fig. S3**, to establish whether recurrent sites were occupied more often than expected by chance, we calculated the probability to observe any given occupancy (including the measured occupancy) using a hypergeometric function.

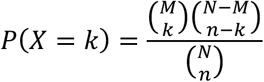

Where, P = the probability of having k recurrent sites occupied, N = potential sites that can be occupied, M = recurrent sites available, n = sites that can be occupied on a given day, k = recurrent sites occupied on a given day. The measured occupancy was significantly different from chance if the sum of all probabilities bigger than the probability observed for the data was greater than 0.95 (probability mass function).

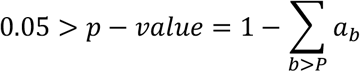

With b = control variable, P = probability and a_b_ = function of control variable.

In Fig.s 2, 3 and S3, to quantify the confidence intervals in the measured fractions of spine gain, spine loss, spine turnover, occupancies and flips, we used a 2-level bootstrapping method, which accounts both for heterogeneity between neurons and for heterogeneity between spines within a neuron. In short, for each group of neurons (ArcTom +, ArcTom -, NexTom +, NexTom -), we randomly sampled (with replacement) the corresponding number of neurons from that group. For each of these sampled neurons, we randomly sampled (with replacement) the corresponding number of spines or sites. Then, we used these sampled spines to compute surrogate spine gain, spine loss, spine turnover, occupancies and flips. This procedure was repeated 10,000 times and the error bars denote the standard deviations of the distribution of these quantities across the different samples. To test the significance of the difference between tdTomato + and tdTomato – groups, we used a permutation test (10^5^ permutations) on the identities of the neurons (ArcTom + vs. ArcTom - or NexTom + vs. NexTom -). To test the significance of a change following EE (ArcTom + before EE vs. ArcTom + after EE *etc*.) we used a paired test: we measured, for each neuron, the difference between the measure (fractions of spine gain, spine loss, spine turnover, occupancies and flips) before and after the EE, and used a two-sided Wilcoxon rank sum test to compute the significance of the change.

