## Supplemental figures for "Stability of excitatory structural connectivity predicts the probability of CA1 pyramidal neurons to become engram neurons"

### SUPPLEMENTARY FIGURES

#### Figure S1. Characterization of Tamoxifen-dependent tdTomato expression.

(**A**) Picture showing a representative Enriched Environment (EE).

(**B**) Confocal images of the dorsal CA1 of Arc-Cre$^{ERT2}$; Ai9 animals after 2h permanence in home cage (HC, upper row) or in EE (lower row). Mice received either 75 mg/kg, 150 mg/kg or 300 mg/kg TAM. Single z-plane (5 µm z-step, 8-10 focal planes). Scale bar, 100 µm

(**C**) Proportion of Arc-tdTomato (ArcTom) + neurons after after HC (black circles) or EE (blue circles). **Table S1** for p values. Bars are S.E.M.

(**D**) A 75 mg/kg TAM injection led to a 2.5 fold increase of ArcTom+ neurons after EE compared to HC.

(**E**) Confocal images of the dorsal CA1 of Nex-Cre$^{ERT2}$; Ai9 animals after 2h permanence in HC (up) or in EE (low). Mice received 75 mg/kg TAM. Single z-plane (5 µm z-step, 8-10 focal planes). Scale bar, 100 µm

(**F**) The proportion of Nex-tdTomato (NexTom) + (empy circles) after 75 mg/kg TAM injection was not different between exposure to HC (black) and EE (blue), differently to ArcTom + neurons (solid circles). Bars are S.E.M. **Table S1** for p values.

(**G**) Schematic definition of the dendritic order of the imaged dendrites.

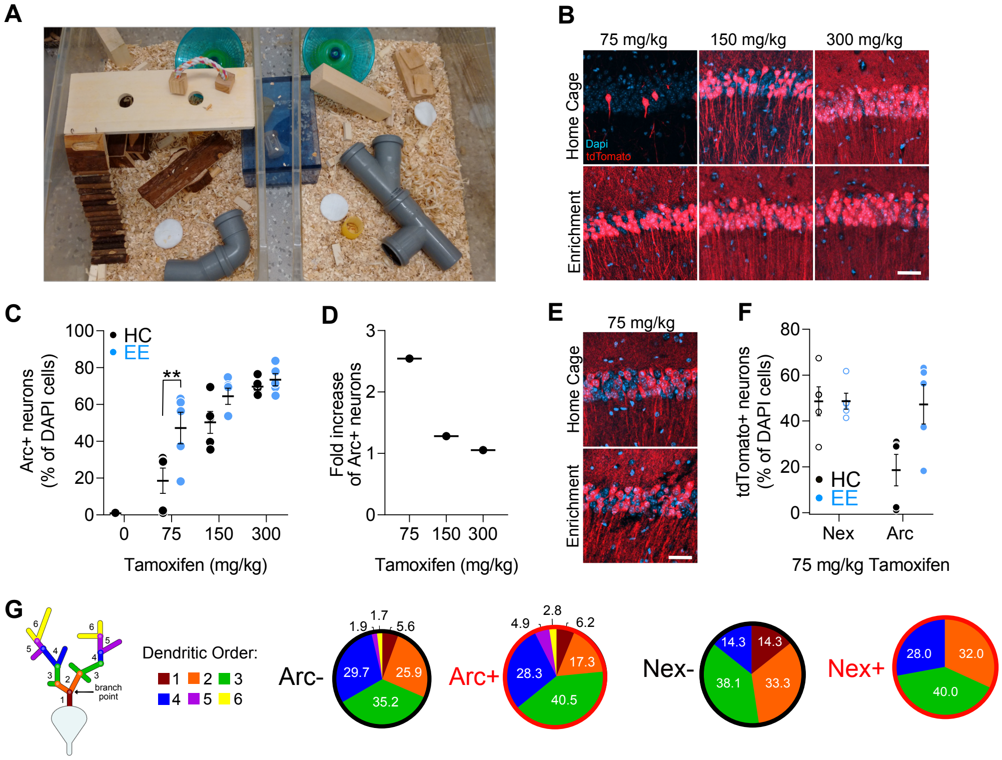

A
B
75 mg/kg
150 mg/kg
300 mg/kg
Home Cage
Enrichment
Dapi
tdTomato
C
HC
EE
**
Arc+ neurons
(% of DAPI cells)
100
80
60
40
20
0
0
75
150
300
Tamoxifen (mg/kg)
D
Fold increase
of Arc+ neurons
3
2
1
0
75
150
300
Tamoxifen (mg/kg)
E
75 mg/kg
Home Cage
Enrichment
F
tdTomato+ neurons
(% of DAPI cells)
80
60
40
20
0
HC
EE
Nex
Arc
75 mg/kg Tamoxifen
G
branch point
Dendritic Order:
1 2 3
4 5 6
Arc-
1.9
1.7
5.6
29.7
25.9
35.2
Arc+
4.9
2.8
6.2
28.3
17.3
40.5
Nex-
14.3
14.3
38.1
33.3
Nex+
28.0
32.0
40.0

**Figure S2. Comparisons of fractional turnover, gain and loss per epoch.**

(**A**) Dendritic spine turnover was higher in prospective ArcTom - than in ArcTom + neurons during baseline, but not after EE. The turnover of ArcTom - neurons decreased after EE.

(**B**) Dendritic spine gain was higher in prospective ArcTom - than in ArcTom + neurons during baseline, but not after EE. The gain of ArcTom - neurons decreased after EE.

(**C**) Dendritic spine loss did not differ between prospective ArcTom - and ArcTom + neurons and was not affected by EE.

(**D** - **F**) Baseline dendritic spine turnover, gain and loss of NexTom - and NexTom + neurons, were not different and were not affected by TAM injection.

(**A** - **F**) Boxes are the second and third quartile while whiskers are the first and last quartiles of the distributions of fractional turnover, gain, and loss of spines per neuron pooled over the epochs reported in the panels. Red: prospective or actual td-Tomato + neurons; grey: prospective or actual td-Tomato - neurons.

(**G**) The spine size of ArcTom - and ArcTom + neurons was stable through time and did not differ between the two groups. Each dot represents the measured size of a persistent spine at each imaging time point. Bars are S.E.M.

**(A - G) Table S1** for p values. Blue vertical line, time point of EE+TAM injection; grey vertical line, time point of TAM injection.

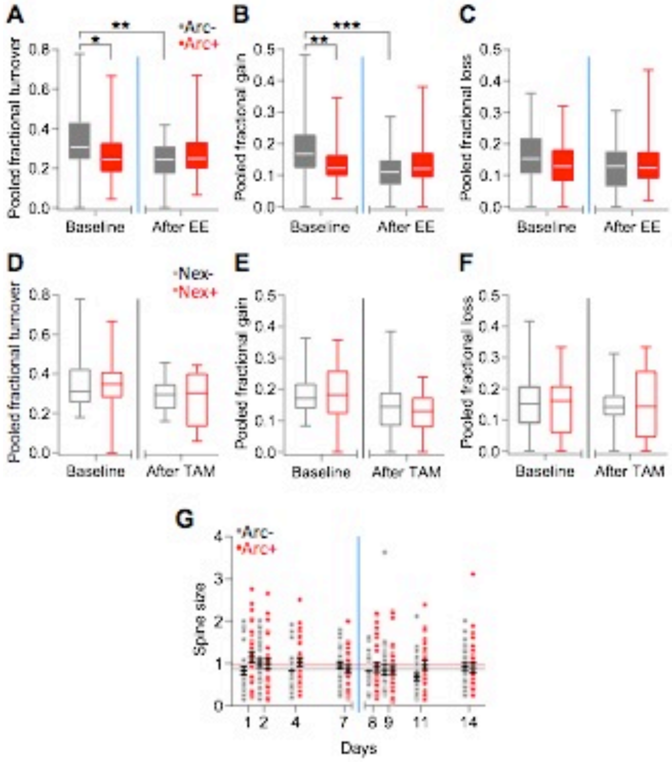

A
B
C
D
E
F
G
Arc-
Arc+
Nex-
Nex+
Pooled fractional turnover
Pooled fractional gain
Pooled fractional loss
Baseline
After EE
After TAM
Spine size
Days

**Figure S3. Occupancy and flips of recurrent sites.**

(**A**, **B**) Recurrent sites of ArcTom + and ArcTom - (solid circles, red and black respectively) and NexTom + and NexTom - (empty circles, red and black respectively) neurons were significantly more occupied than expected by chance on most time points.

(**C - J**) Histograms of the number of time point occupied (C - F) or the number of flips (G - J) of recurrent synaptic sites of prospective ArcTom + (full, red) , NexTom + (empty, red) and ArcTom - (full, black) and NexTom - (empty, black) neurons during baseline (C, E, G, I), after EE (D, H) or TAM injection (I, J). **S.T.A.R. methods** for defintion of the error bars.

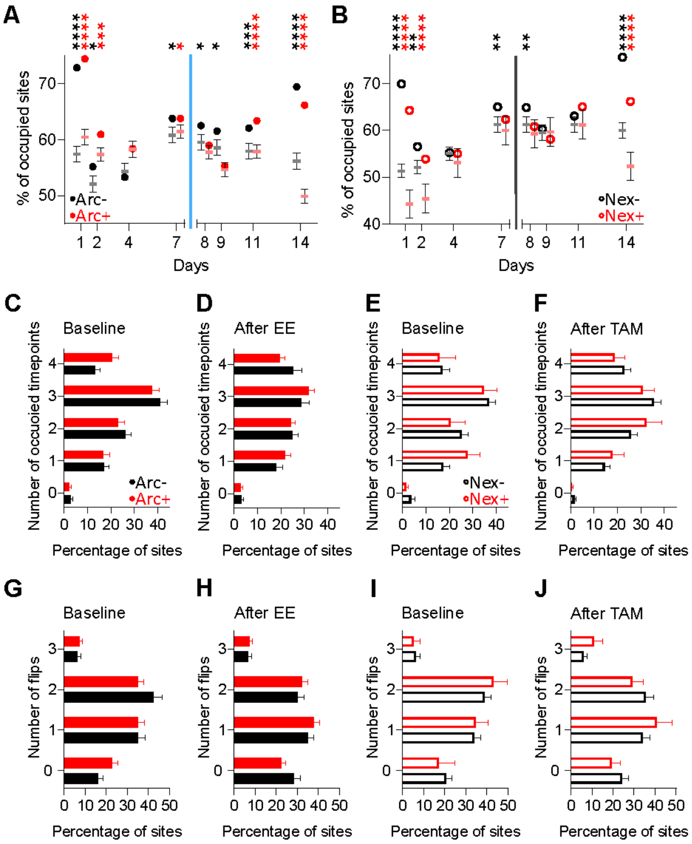

A
B
C
D
E
F
G
H
I
J
% of occupied sites
Days
Arc-
Arc+
Nex-
Nex+
Baseline
After EE
After TAM
Number of occuoied timepoints
Number of flips
Percentage of sites

**Figure S4. After the EE or TAM the density and survival of dendritic spines of CA1 pyramidal neurons does not correlate with freezing in a trace fear conditioning task.**

(**A** - **D**) The density of spines after EE+TAM injection or after TAM injection of Arc-Cre$^{ERT2}$; Ai9; Thy1-eGFP (solid circles) and Nex-Cre$^{ERT2}$; Ai9; Thy1-eGFP (open circles) mice did not correlate to freezing to the context or to the tone.

(**E** - **H**) The time constant of the surviving fraction of spines after EE+TAM injection or after TAM injection of Arc-Cre$^{ERT2}$; Ai9; Thy1-eGFP; (solid circles) and Nex-Cre$^{ERT2}$; Ai9; Thy1-eGFP (open circles) mice did not correlate to freezing to the context or to the tone.

(**I** - **L**). The time constants of the fraction gained of spines after the EE+TAM injection or after TAM injection of Arc-Cre$^{ERT2}$; Ai9; Thy1-eGFP (solid circles) and Nex-Cre$^{ERT2}$; Ai9; Thy1-eGFP (open circles) mice did not correlate to freezing to the context or to the tone.

**Table S1** for p values.

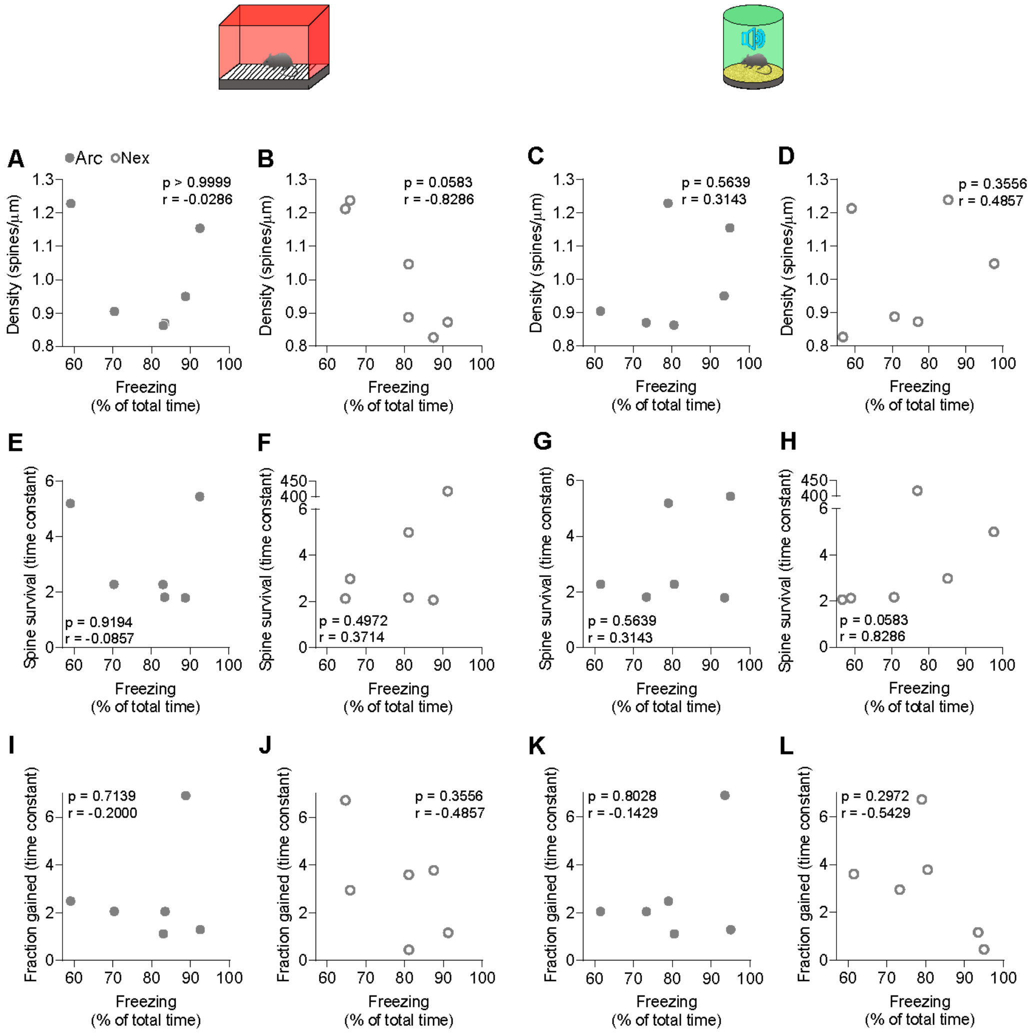

A
Arc
Nex
p > 0.9999
r = -0.0286
B
p = 0.0583
r = -0.8286
C
p = 0.5639
r = 0.3143
D
p = 0.3556
r = 0.4857
Density (spines/µm)
E
p = 0.9194
r = -0.0857
F
p = 0.4972
r = 0.3714
G
p = 0.5639
r = 0.3143
H
p = 0.0583
r = 0.8286
Spine survival (time constant)
I
p = 0.7139
r = -0.2000
J
p = 0.3556
r = -0.4857
K
p = 0.8028
r = -0.1429
L
p = 0.2972
r = -0.5429
Fraction gained (time constant)
Freezing
(% of total time)
